# Gamma-frequency firing persistently enhances intrinsic excitability in retinal ganglion cells

**DOI:** 10.64898/2026.10.22.760873

**Authors:** Prithvi Ramtohul, Laure Molinieres, Victor Calbiague-Garcia, Maria Laura Musella, Fabrice Abate, Aurore Aziz, Thierry David, Danièle Denis, Olivier Marre, Michaël Russier, Dominique Debanne

## Abstract

Experience-dependent plasticity is usually attributed to synaptic changes, but neuronal output also depends on changes in intrinsic excitability. Whether intrinsic excitability is persistently modified at the retinal output stage is unknown. Using whole-cell recordings from retinal ganglion cells (RGCs) of juvenile rats, we found that 40-Hz firing produced sustained potentiation of intrinsic excitability in low-input-resistance A-type RGCs and rapid potentiation of intrinsic excitability in high-input-resistance B-type RGCs. In A-type bursting RGCs, long-term potentiation of intrinsic excitability (LTP-IE) required Cav1 calcium channels for its induction. LTP-IE was associated with hyperpolarization of spike threshold and reduction of spike time jitter, due to reduced Kv1 potassium-family channel function. Potentiation also increased the probability that visually-evoked synaptic events triggered action potentials. Patterned firing therefore persistently reconfigures RGC input-output transformation through the Cav1-dependent regulation of Kv1 channels, revealing intrinsic plasticity at the retinal output stage.

## Introduction

Visual experience shapes the postnatal maturation of visual function (Hooks & Chen, 2020). Disruption of patterned vision during critical periods can produce enduring deficits, most notably reduced visual acuity (Morishita & Hensch, 2008), implicating activity-dependent plasticity in the developmental improvement of vision. Such plasticity has been characterized extensively in the visual cortex after monocular deprivation (Wiesel & Hubel, 1963), rearing in orientation-restricted environments (Blakemore & Cooper, 1970), or cellular conditioning (Frégnac *et al*., 1988; Debanne *et al*., 1998; Schuett *et al*., 2001). More recently, experience-dependent plasticity has also been demonstrated in visual thalamus after monocular deprivation (Jaepel *et al*., 2017; Sommeijer *et al*., 2017; Duménieu *et al*., 2025), indicating that adaptive changes occur at multiple levels of the visual pathway.

Evidence that visual experience also reshapes retinal circuits has accumulated. Retinal ganglion cells (RGCs), the output neurons of the retina, integrate signals from retinal interneurons and transmit visual information to the brain through the optic nerve (Sanes & Masland, 2015). Visual activity is required for postnatal refinement of ON and OFF pathways (Feller, 2003; Tian & Copenhagen, 2003) and can modify glutamate-receptor composition in ON RGCs (Xia *et al*., 2007). Early visual deprivation alters RGC receptive fields (Di Marco *et al*., 2009) and optic-nerve conduction (Etxeberria *et al*., 2016), whereas dark rearing changes GABAergic transmission in rod bipolar cells (Wisner *et al*., 2023). Genetic suppression of the retinal ON pathway also reduces the intrinsic excitability of ON RGCs (Khoussine *et al*., 2025), and patterned visual experience modifies retinal cellular morphology and functional output in zebrafish RGCs (Reynolds *et al*., 2026). Together, these observations suggest that retinal output can adapt to sustained changes in visual activity.

Long-lasting plasticity in visual circuits has classically been attributed to persistent changes in synaptic transmission (Kleinschmidt *et al*., 1987; Law & Cooper, 1994). However, neuronal input-output transformations can also be modified through plasticity of intrinsic excitability (IE) (Debanne *et al*., 2019). Long-term intrinsic plasticity has been described in visual cortical neurons (Cudmore & Turrigiano, 2004; Nataraj *et al*., 2010) and in relay neurons of the dorsal lateral geniculate nucleus (Duménieu *et al*., 2025; Aziz *et al*., 2026), but whether RGCs express a comparable form of activity-dependent intrinsic plasticity has not been tested directly. Several inhibitory ion channels can be persistently regulated by neuronal activity, including hyperpolarization-activated cyclic nucleotide-gated (HCN) channels (Fan *et al*., 2005; Brager & Johnston, 2007; Campanac *et al*., 2008), Kv1 channels (Campanac *et al*., 2013; Duménieu *et al*., 2025), and Kv7 channels (Incontro *et al*., 2021; Sammari *et al*., 2022). Kv1 channels are particularly well positioned to control spike initiation and timing because they are enriched at the axon initial segment (AIS) (Goldberg *et al*., 2008; Lorincz & Nusser, 2008), and Kv1-family channels are expressed in RGCs (Koeberle *et al*., 2010; Ihadadene *et al*., 2025).

Here, we asked whether patterned spiking can induce persistent intrinsic plasticity in RGCs. We show that 40-Hz firing produces a slowly developing long-term potentiation of intrinsic excitability (LTP-IE) in low-input-resistance A-type RGCs and a rapid potentiation in high-input-resistance B-type RGCs. In A-type bursting RGCs, LTP-IE requires Cav1 channels for induction and is associated with reduced Kv1-family channel function, leading to a more hyperpolarized action-potential threshold, shorter spike latency, and greater spike-time precision. This intrinsic plasticity also increases the probability that visually evoked synaptic input triggers action potentials, demonstrating that activity-dependent changes in intrinsic excitability can reshape retinal output.

## Results

### Gamma-frequency firing induces intrinsic plasticity in RGCs

RGCs were recorded in whole-cell current clamp in flat-mounted retinas (**Figure 1A**). Intrinsic properties were measured in the presence of the synaptic blockers kynurenate and picrotoxin. Following the physiological classification of O’Brien et al. (O’Brien *et al*., 2002), cells were separated by input resistance (Rin) using an 800-MΩ criterion (**Figure 1B, C**). Rin was 461 ± 34 MΩ in A-type RGCs (n = 10) and 1553 ± 170 MΩ in B-type RGCs (n = 5) (**Figure 1C**). Non-bursting and bursting subtypes were identified from the presence or absence of a rebound burst after hyperpolarization (Margolis & Detwiler, 2007; Werginz *et al*., 2025). ON / OFF classification of RGC was made in a few cases with visual stimuli. In this case, RGCs were recorded without synaptic blockers at physiological external calcium and magnesium concentrations (1.5 and 1.0 mM, respectively; **Figure S1**). We then tested whether both RGC A et B classes expressed activity-dependent intrinsic plasticity using a protocol that induces LTP-IE in visual cortical and thalamic neurons (Cudmore & Turrigiano, 2004; Nataraj *et al*., 2010; Russier *et al*., 2024; Duménieu *et al*., 2025). Trains of 15 action potentials (APs) at 40 Hz were evoked at 0.1 Hz for 10 min (**Figure 1D**). In A-type RGCs, excitability increased gradually after the induction protocol (**Figure 1E**). Between 20 and 30 min after induction, AP number reached 146 ± 11% of baseline (n = 10), increasing from 6.0 ± 0.9 to 8.7 ± 1.2 APs (**Figure 1E**), without a significant change in input resistance (**Figure S2A, B**). By contrast, B-type RGCs showed a rapid increase in excitability after 40-Hz firing (**Figure 1F, G**): AP number increased from 3.8 ± 0.6 to 5.3 ± 0.8 (n = 5) within the first 2 min after stimulation (**Figure 1G**). To distinguish this rapid early response from the slowly developing LTP-IE in A-type RGCs, we operationally define the potentiation measured 2 min after induction as short-term potentiation of intrinsic excitability (STP-IE; **Figure S2D**). Again, no detectable change in input resistance was observed (**Figure S2C**). Potentiation at 2 min was greater in B-type than in A-type RGCs (**Figure S2D**). Thus, patterned firing produces intrinsic plasticity with distinct kinetics in low- and high-input-resistance RGCs, expressed as slowly developing LTP-IE in A-type RGCs and immediate potentiation of IE in B-type RGCs.

**Figure 1.**
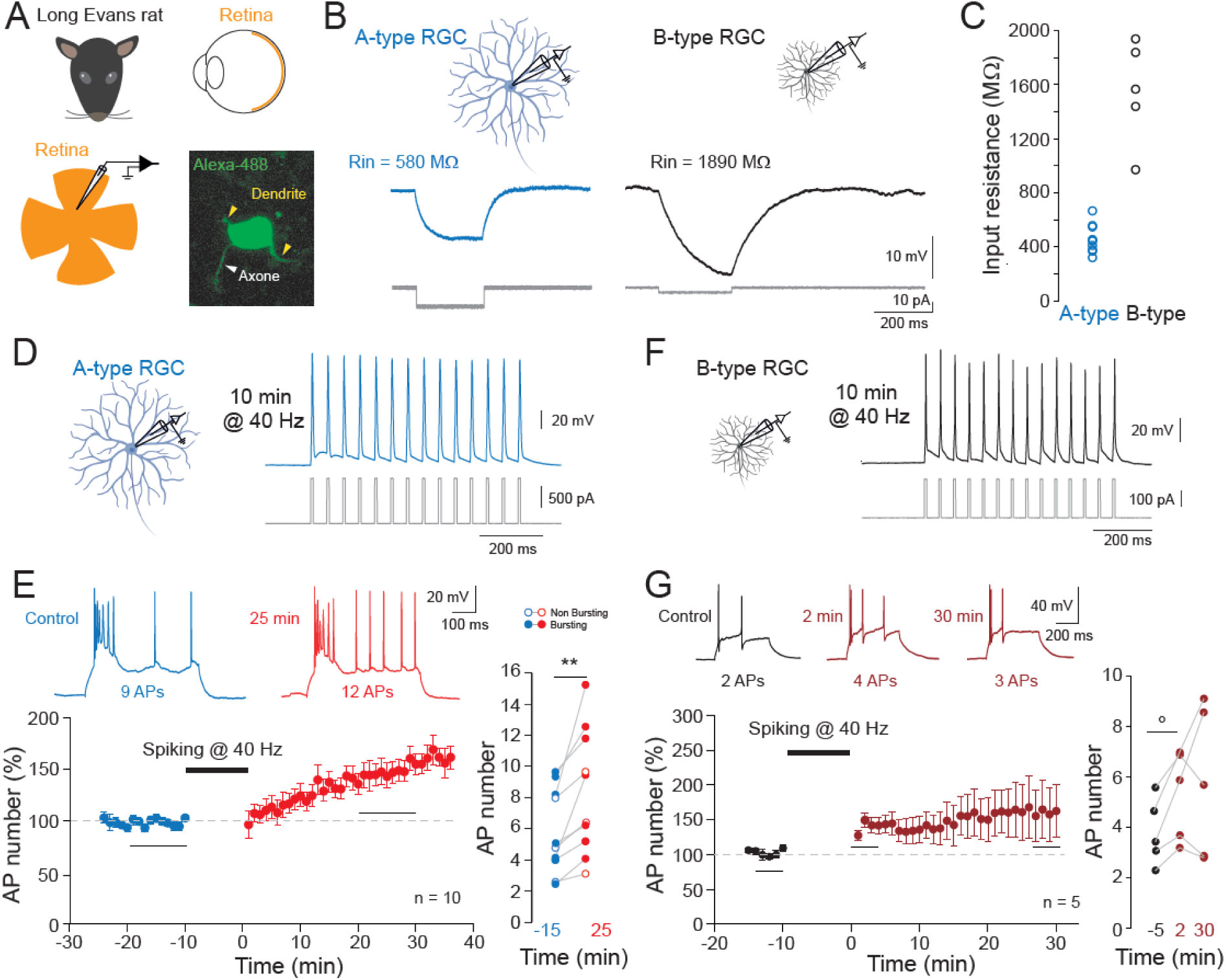
Induction of intrinsic plasticity in RGCs. (A) Experimental preparation. The eye was isolated from a P17-P23 Long Evans rat (top), and the retina was flat-mounted for patch-clamp recording. Bottom right, confocal image of an RGC filled with Alexa Fluor 488, showing the soma, axon, and dendritic arbor. (B) Representative membrane responses of a low-input-resistance A-type RGC (left) and a high-input-resistance B-type RGC (right). (C) Distribution of input resistance in A- and B-type RGCs. (D) Forty-hertz induction protocol in an A-type RGC. (E) LTP-IE in A-type RGCs. Left, time course of normalized AP number; top, representative traces before and 25 min after induction. Right, paired AP counts before and after induction. **, Wilcoxon signed-rank test, p < 0.01. (F) Forty-hertz induction protocol in a B-type RGC. (G) Rapid intrinsic potentiation in B-type RGCs. Left, time course; top, representative traces at baseline, 2 min, and 30 min after induction. Right, paired AP counts. °, Wilcoxon signed-rank test, p < 0.07. RGC schematics were created with BioRender.

### Cav1 channels are required for LTP-IE induction in A-type bursting RGCs

We next examined the induction mechanism of LTP-IE in A-type bursting RGCs. A similar 40-Hz protocol in visual thalamic neurons requires Cav1 (L-type) calcium channels activated by spike depolarization (Duménieu *et al*., 2025), and Cav1 channels are expressed in RGCs (Van Hook *et al*., 2019). We therefore applied nitrendipine (25 µM), a Cav1 channel blocker, during the induction protocol. Under Cav1 blockade, 40-Hz firing did not produce the sustained potentiation observed in control cells; AP number reached 98 ± 6% of baseline (3.9 ± 0.3 AP in control vs. 3.9 ± 0.5 AP after 40 Hz stimulation, n = 8; **Figure 2A**). The normalized AP changes were statistically different from control data (**Figure 2B**). These results indicate that Cav1-channel activation is required for LTP-IE induction in A-type bursting RGCs (**Figure 2C**).

**Figure 2.**
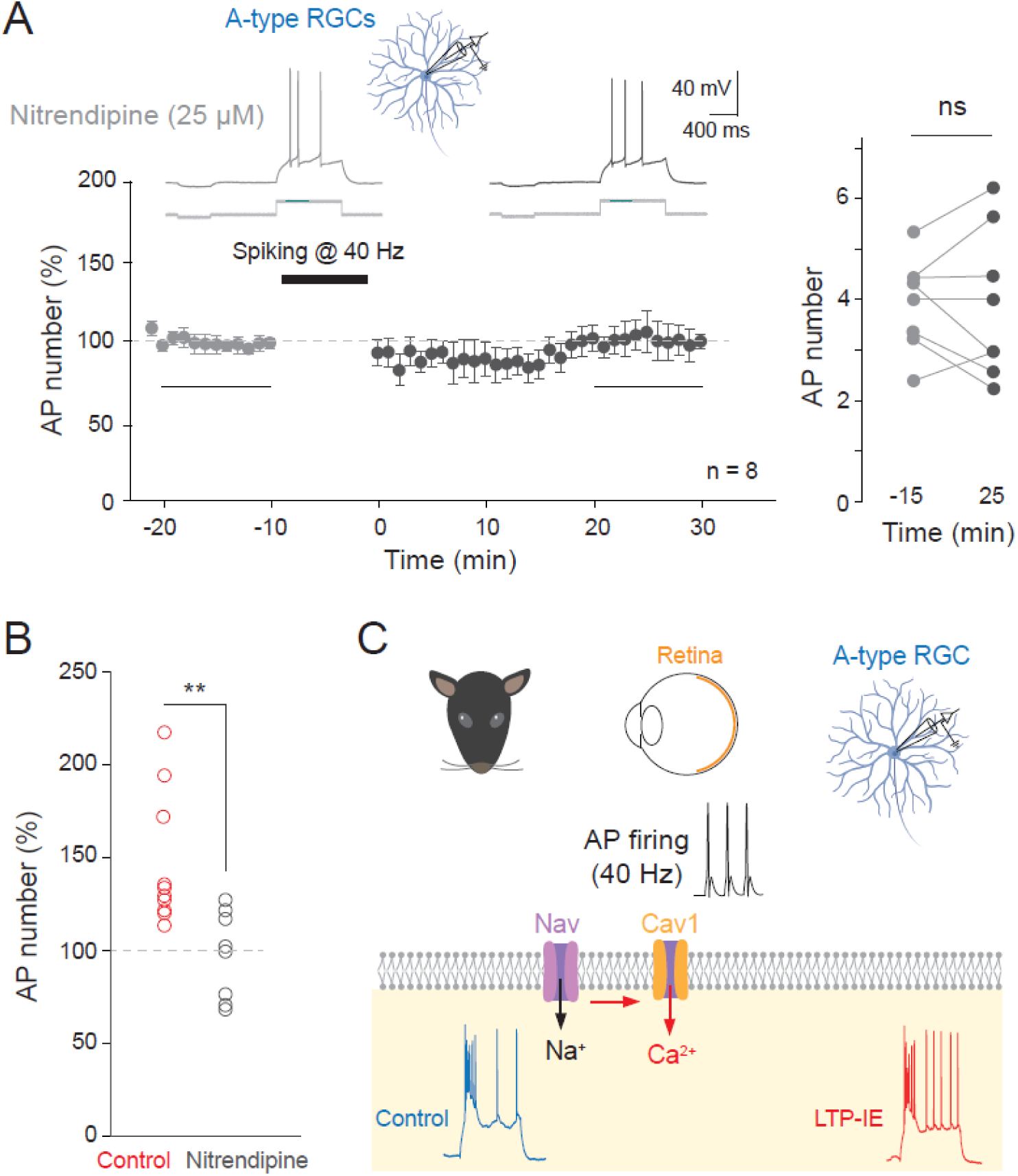
Cav1 (L-type) calcium channel blockade prevents LTP-IE induction in A-type bursting RGCs. (A) Time course of normalized AP number during bath application of nitrendipine (25 µM) in A-type bursting RGCs. The 40-Hz induction protocol was delivered at time -10 (for 10 min). Right, paired AP counts before and 25 min after induction (n = 8). ns, Wilcoxon, p > 0.1. (B) Comparison of the normalized changes in control and in nitrendipine. **, Mann-Whitney Y-test, p < 0.01. (C) Working model for LTP-IE induction: AP firing activates Nav channels and Cav1 channels, and Ca^2+^ entry through Cav1 channels is required for the induction of persistent intrinsic potentiation. Retina and RGC schematics were created with BioRender.

### Reduced Kv1-family channel function mediates LTP-IE expression

To identify mechanisms underlying LTP-IE expression, we first measured AP threshold before and after induction. A-type bursting RGCs displayed a T-type calcium-dependent rebound burst (**Figure S3**) followed by regular spiking, allowing us to measure separately the threshold of the first AP in the burst and that of the first regular AP (**Figure 3A**). After LTP-IE induction, the threshold of the first burst AP was hyperpolarized by ∼2.2 mV (**Figure 3B**), and the threshold of the first regular AP was hyperpolarized by ∼2.5 mV (**Figure 3C**). Thus, increased intrinsic excitability was accompanied by a shift in spike threshold toward more negative potentials.

**Figure 3.**
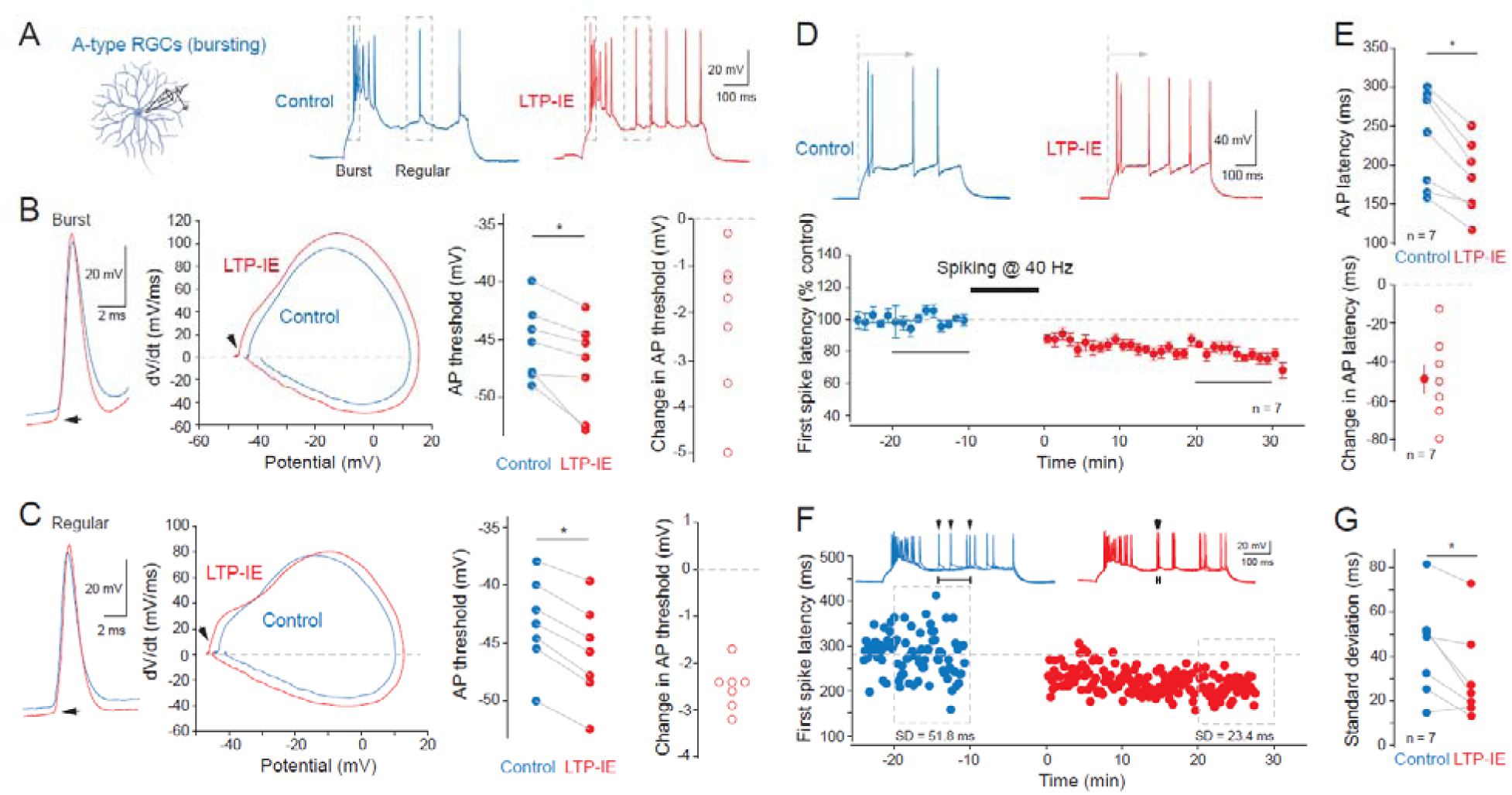
LTP-IE hyperpolarizes AP threshold and improves spike timing. (A) Recording configuration and representative voltage traces from an A-type bursting RGC showing the initial rebound burst and the first regular AP before and after LTP-IE induction. (B) Threshold of the first AP in the rebound burst. Left, representative voltage traces and phase plots; middle right, paired thresholds; right, change in threshold. *, Wilcoxon signed-rank test, p < 0.05. (C) Threshold of the first regular AP, displayed as in (B). *, Wilcoxon signed-rank test, p < 0.05. (D) Representative traces and time course of the first regular AP latency before and after 40-Hz induction. (E) Group data for absolute latency (top) and change in latency (bottom). (F) Representative reduction in first-spike latency variability after LTP-IE; the standard deviation decreased from 51.8 to 23.4 ms in the example shown. (G) Group data for spike-time variability. *, Wilcoxon signed-rank test, p < 0.05. RGC schematic created with BioRender.

LTP-IE also altered spike timing. The latency of the first regular AP decreased by ∼50 ms after induction (**Figure 3D, E**), and trial-to-trial variability in first-spike latency was reduced (**Figure 3F, G**). The combination of threshold hyperpolarization, shorter latency, and reduced spike-time jitter is consistent with decreased Kv1-family channel activity (Goldberg *et al*., 2008; Cudmore *et al*., 2010; Campanac *et al*., 2013; Duménieu *et al*., 2025).

We therefore asked whether pharmacological Kv1 blockade would mimic and occlude LTP-IE. Dendrotoxin-I (DTx-I), which blocks Kv1.1-, Kv1.2-, and Kv1.6-containing channels, reproduced the principal features of LTP-IE: AP number increased, AP threshold became more hyperpolarized, and first-spike latency decreased (**Figure 4A**). Moreover, when Kv1 channels were blocked with DTx-I, the 40-Hz protocol produced no further sustained potentiation (3.2 ± 0.6 AP in control vs. 3.5 ± 0.7 AP after 40 Hz stimulation corresponding to 112 ± 5% of baseline AP number, n = 7; **Figure 4B**), consistent with occlusion. In addition, the Together, these results support a model in which Cav1-dependent calcium entry induces LTP-IE through a persistent reduction in Kv1-family channel function (**Figure 4C**).

**Figure 4.**
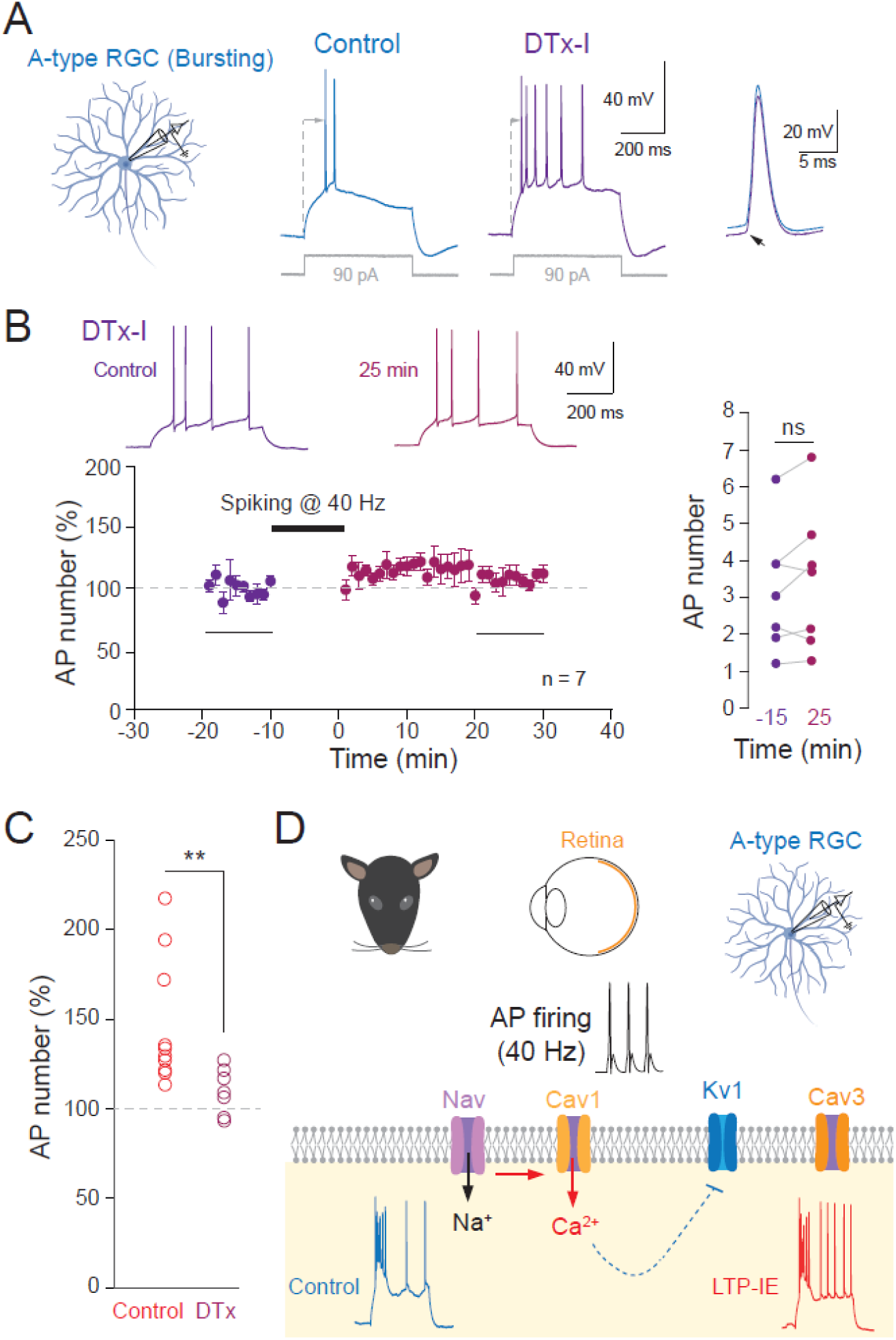
Kv1 channel blockade mimics and occludes LTP-IE in RGCs. (A) DTx-I inhibition of Kv1-family channels mimics key features of LTP-IE in A-type bursting RGCs, increasing AP number and hyperpolarizing AP threshold. (B) LTP-IE is occluded in the presence of DTx-I. Left, time course; right, paired AP counts before and 25 min after 40-Hz firing. ns, Wilcoxon signed-rank test, p > 0.1. (C) Comparison of the normalized changes in AP number in control and in DTx. **, Mann-Whitney U-test, p < 0.01. (C) Working model: 40-Hz firing recruits Cav1-dependent Ca^2+^ entry, which leads indirectly to a persistent reduction in Kv1-family channel function and increased intrinsic excitability.

### LTP-IE enhances spike generation in visually-evoked responses

The preceding experiments quantified intrinsic excitability with somatic current injection, leaving open whether LTP-IE also changes the conversion of synaptic input into spikes. To test this, we next determined whether visually-evoked synaptic responses could carry LTP-IE in A-type bursting RGCs. Wide field visual stimulation was used to assess RGC excitability before and after 40 Hz spiking activity. These experiments were performed in physiological external Ca^2+^ and Mg^2+^ concentration, (1.5 and 1.0 mM, respectively). A significant enhancement of spike number was observed in bursting A-type RGCs following spiking at 40 Hz (**Figure 5A**). No change in input resistance was found (**Figure 5B**). In a manner similar to what has been observed with pulses, the AP threshold was hyperpolarized by ∼5 mV (**Figure 5C**) and the delay to the first spike was reduced by ∼20 ms (**Figure 5D**). Similar findings were also observed in another ON RGC (**Figure S4**). We conclude that visually-evoked synaptic responses provide a functional readout of LTP-IE.

**Figure 5.**
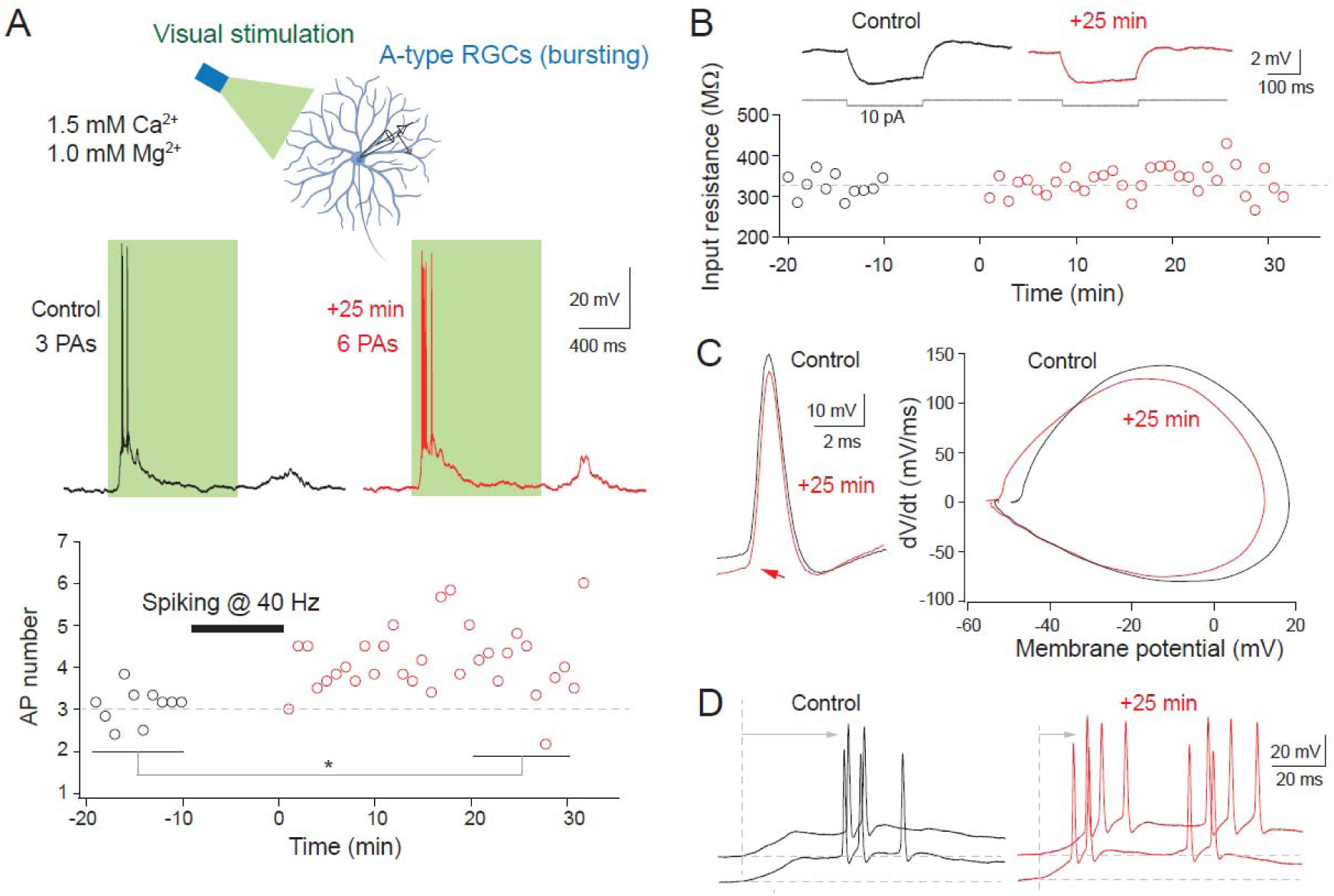
LTP-IE in visually-evoked synaptic responses. (A) Lasting enhancement of spiking activity evoked by green light in a A-type bursting RGC. Top, scheme of the recording configuration. Middle, representative voltage traces before (black) and after (red) spiking activity at 40 Hz showing LTP-IE of visually evoked responses. Bottom, time course of AP number before and after 40 Hz firing. *, Wilcoxon test, p<0.05. (B) Stability of input resistance. Top, traces. Bottom, time course. (C) Hyperpolarized AP threshold. Left, visually evoked first APs before (black) and after LTP-IE induction (red). Right, phase plots showing an ∼5 mV hyperpolarization of AP threshold after LTP-IE induction. (D) Shortening of the AP delay after LTP-IE. RGC schematic created with BioRender.

## Discussion

We identify a persistent, activity-dependent increase in intrinsic excitability at the retinal output stage. Ten minutes of 40-Hz firing produced slowly developing LTP-IE in low-input-resistance A-type RGCs and a rapid potentiation in high-input-resistance B-type RGCs. In A-type bursting RGCs, LTP-IE required Cav1 channels for induction and was associated with a more hyperpolarized AP threshold, shorter first-spike latency, and reduced spike-time variability. Pharmacological blockade of Kv1 channels reproduced these excitability changes and occluded further LTP-IE, consistent with a mechanism in which Cav1-dependent calcium entry leads to a persistent reduction in Kv1 channel activity. Importantly, LTP-IE also increased the probability that visually-evoked synaptic activity triggered APs, linking intrinsic plasticity to a functional enhancement of retinal output.

### Gamma-frequency firing as an inducer of RGC intrinsic plasticity

LTP-IE was induced by brief episodes of repetitive firing at 40 Hz for 10 min, a protocol previously used to induce intrinsic plasticity in layer 5 visual cortical neurons (Nataraj *et al*., 2010) and dLGN relay neurons (Russier *et al*., 2024; Duménieu *et al*., 2025; Aziz *et al*., 2026). This firing rate lies within the physiological range of RGC activity, which can transiently reach 50 to 80 Hz (Kuffler, 1953), and gamma-frequency retinal oscillations have been described in cat and mouse retina (Neuenschwander & Singer, 1996; Storchi *et al*., 2017). We did not anatomically classify the recorded cells, so correspondence with canonical morphological RGC classes remains provisional. Their passive properties are consistent with low-input-resistance A-type cells representing large, α-like RGCs and high-input-resistance B-type cells representing smaller, β-like populations (O’Brien *et al*., 2002; Sanes & Masland, 2015). The distinct temporal profiles of plasticity in the two physiological classes suggest cell-type-specific mechanisms; the mechanism underlying the rapid potentiation in B-type RGCs remains to be determined.

### Induction and expression mechanisms of LTP-IE in A-type RGCs

The data support a sequential mechanism in which repetitive AP firing activates voltage-gated sodium channels and, through spike depolarization, recruits Cav1 channels. The resulting calcium entry initiates a persistent reduction in Kv1-family channel function. Three observations support this model. First, nitrendipine prevented the sustained potentiation normally evoked by the 40-Hz protocol. Second, LTP-IE shifted AP threshold to more negative potentials and reduced both first-spike latency and its variability, a signature consistent with reduced Kv1-mediated current (Cudmore *et al*., 2010; Campanac *et al*., 2013; Duménieu *et al*., 2025). Third, DTx-I reproduced these electrophysiological changes and occluded further potentiation, placing Kv1-family channels downstream of Cav1-dependent induction.

RGCs express several Kv1 subunits, including Kv1.1, Kv1.2, Kv1.3, and Kv1.6 (Koeberle *et al*., 2010; Ihadadene *et al*., 2025). Our pharmacology does not resolve which subunit or subunit combination is responsible for LTP-IE. More broadly, intrinsic excitability in RGCs can be modified by other perturbations; for example, early experimental glaucoma increases RGC excitability through Nav1.6 sodium channel up-regulation (Risner *et al*., 2018). In many central neurons, activity-dependent intrinsic plasticity is accompanied by changes in AIS position, length, or ion-channel content (Grubb & Burrone, 2010; Kuba *et al*., 2010, 2015; Fékété *et al*., 2021; Zbili *et al*., 2021; Fréal *et al*., 2023; Sanz-Gálvez *et al*., 2025; Benoit *et al*., 2026; Aziz *et al*., 2026). Because the AIS was not examined here, whether RGC LTP-IE involves altered Kv1 trafficking or abundance, post-translational regulation, or another mechanism remains unresolved.

### Consequences for spike timing and retinal output

LTP-IE increased not only spike number but the delay of spiking was globally reduced (Cudmore *et al*., 2010) while the temporal precision of spike initiation was enhanced. Reduced first-spike latency is expected when Kv1-dependent restraint near threshold is weakened (Higgs & Spain, 2011; Campanac *et al*., 2013), and the accompanying reduction in latency variability should sharpen the temporal fidelity of RGC output. Axonal sodium-channel availability and propagation can influence how somatic timing changes are transmitted downstream RGCs (Zbili & Debanne, 2020; Zbili *et al*., 2020). More generally, retinal and cortical visual coding can depend on relative spike latency and millisecond-scale timing (Celebrini *et al*., 1993; Thorpe *et al*., 1996, 2001; Gollisch & Meister, 2008). Thus, intrinsic plasticity has the potential to modify not only the gain of retinal output but also its temporal code, although the consequences for visually evoked signaling remain to be established directly.

### Visually-evoked synaptic responses provide a functional readout of LTP-IE

We show here that visually-evoked synaptic responses carry LTP-IE in A-type bursting RGCs in a similar way to current pulses. The AP threshold was hyperpolarized and the delay to the first spike was reduced by ∼20 ms. Although this observation has been made on only a few RGCs, it strongly suggests that visually-evoked synaptic responses provide a functional readout of LTP-IE.

### Intrinsic plasticity broadens the mechanisms of retinal adaptation

Retinal adaptation is increasingly recognized to involve multiple forms of plasticity. OFF transient α-RGCs, for example, show pronounced short-term depression of visually evoked excitatory transmission (Chen & Rieke, 2026), and dark rearing disrupts the normal developmental segregation of ON and OFF pathways (Tian & Copenhagen, 2003). Our results add a cell-intrinsic component to these synaptic mechanisms: patterned spiking can persistently alter the input-output function of RGCs through Cav1- and Kv1-dependent regulation of excitability. Because LTP-IE was induced here by current-evoked firing, a key next step will be to determine whether defined visual stimulation—particularly gamma-range stimulation—can induce the same plasticity under physiological conditions and whether it measurably changes visual coding. Intrinsic excitability should therefore be considered alongside synaptic plasticity as a potential substrate of experience-dependent adaptation in the retina (Baden *et al*., 2016).

## Materials and Methods

### Ex vivo retinal preparation

Experimental procedures were approved by the Préfecture des Bouches-du-Rhône (DDPP13, approval number: # E 13-055-08). P17-P23 Long Evans rats were deeply anesthetized with 5% isoflurane and euthanized by decapitation. Eyes were rapidly enucleated and placed in oxygenated extracellular solution containing (in mM) 125 NaCl, 2.5 KCl, 3 CaCl_2_, 2 MgCl_2_, 1.09 NaH_2_PO_4_, 26 NaHCO_3_, and 10 D-glucose, equilibrated with 95% O_2_/5% CO_2_ (pH 7.4). Under red illumination, the cornea, iris, and lens were removed, the retina was gently separated from the eyecup, and the vitreous was removed with forceps (Calbiague *et al*., 2020; Goethals *et al*., 2025). Retinas were then incubated for 10-15 min in extracellular solution containing collagenase CLS-1 (3.2 mg in 5 ml; Serlabo) to digest residual vitreous. After thorough rinsing, retinas were kept in the dark in oxygenated extracellular solution.

For visual stimulation, retinas were dissected in an extracellular solution containing (in mM) 125 NaCl, 2.5 KCl, 1.5 CaCl_2_, 1 MgCl_2_, 1.09 NaH_2_PO_4_, 26 NaHCO_3_, 20 D-glucose, and 0.431 L-glutamine, with 95% O_2_/5% CO_2_ (pH 7.4). In this case, retinas were not incubated in collagenase solution.

A retinal piece was transferred to the recording chamber with the ganglion-cell layer facing upward and secured with a small platinum grid.

### Patch-clamp recordings

RGCs were identified by their round somata near the retinal surface. The recording solution contained (in mM) 125 NaCl, 26 NaHCO_3_, 3 CaCl_2_, 2.5 KCl, 2 MgCl_2_, 1.09 NaH_2_PO_4_, and 10 D-glucose and was equilibrated with 95% O_2_/5% CO_2_. GABAergic inhibition was blocked with picrotoxin (100 µM), and excitatory synaptic transmission was blocked with kynurenate (2 mM), unless otherwise stated. Patch pipettes (5-10 MΩ) were pulled from borosilicate glass with a Narishige PC-100 puller and filled with intracellular solution containing (in mM) 120 K-gluconate, 20 KCl, 10 HEPES, 0.5 EGTA, 2 MgCl_2_, 2 Na_2_ATP, and 0.3 NaGTP (pH 7.4). Recordings were performed at 30°C with a MultiClamp 700B amplifier (Molecular Devices) in a temperature-controlled chamber (Luigs & Neumann, Ratingen, Germany). Membrane potentials were not corrected for the -13-mV liquid junction potential. Access resistance was compensated using bridge balance and capacitance neutralization (>70%). Cells were recorded in current clamp, and input resistance was monitored throughout each experiment; recordings in which input resistance varied by >20% were excluded. Voltage and current signals were low-pass filtered at 10 kHz and sampled at 20 kHz with pClamp (Molecular Devices). Standard acquisition epochs lasted 2 s; visual-stimulation recordings lasted 20 s.

### Visual stimulation

To classify RGCs into ON or OFF cells, ganglion cells were visually stimulated for 1-2 sec with a computer-controlled illumination system (CoolLED pE-100) through the microscope. For this purpose, RGCs were recorded in external solution containing physiological Ca²⁺ (1.5 mM) and Mg²⁺ (1.0 mM) and visual stimulation consisted in a full-field green light (λ = 530 nm) delivered for 1-2 s at 0.1 Hz using a triggered illumination system (CoolLED pE-100) set to 0.1% of maximal LED intensity (i.e., ∼5-20 µW/cm^2^), which corresponds to a range between 3.82 x 10^4^ R* and 1.53 x 10^5^ R* (iso/ph/s).

### Live-cell imaging

RGC morphology was visualized with Alexa Fluor 488 added to the intracellular solution (50 µM). Cells were maintained in whole-cell configuration for at least 15 min before imaging. Alexa Fluor 488 was excited at 488 nm and imaged with a Zeiss LSM 710 confocal microscope. Z-stacks were acquired at 1-µm intervals and reconstructed in Fiji/ImageJ.

### Induction and quantification of LTP-IE

LTP-IE was induced by trains of 15 APs at 40 Hz, delivered at 0.1 Hz for 10 min (Cudmore & Turrigiano, 2004; Russier *et al*., 2024; Duménieu *et al*., 2025). Intrinsic excitability was monitored with 800-ms depolarizing current steps (150-250 pA) delivered at 0.1 Hz and adjusted to evoke 3-4 APs under baseline conditions. The current-step amplitude was then held constant before and after the induction protocol. LTP-IE was quantified 20-30 min after induction.

### Drugs

All drugs were bath applied using a peristaltic pump. To facilitate solution exchange, the immersion objective was raised from contact with the bath during drug application. Kynurenate (2 mM) was obtained from Sigma-Aldrich, picrotoxin (100 µM) from Abcam, DTx-I (100 nM) from Alomone Labs, and nitrendipine (25 µM) from Tocris Bioscience.

### Statistics

Data are reported as mean ± SEM. Paired comparisons were performed with the Wilcoxon signed-rank test, and comparisons between independent groups were performed with the Mann-Whitney U-test. Statistical significance was defined as p < 0.05. Analyses were performed with the BiostaTGV online statistical platform.

## Acknowledgments

We thank K. Milton and A. Venture for excellent animal care.

## Author contributions

Prithvi Ramtohul performed experiments, analyzed data, and drafted the manuscript; Laure Molinieres performed experiments and conducted data and statistical analyses; Victor Calbiague performed experiments and analyzed data; Maria Laura Musella and Fabrice Abate performed experiments and analyzed data; Aurore Aziz, Danièle Denis and Thierry David contributed to data analysis; Olivier Marre analyzed data and drafted the manuscript; Michaël Russier designed the study, performed experiments and contributed to data and statistical analyses; Dominique Debanne conceived and designed the study, secured funding, analyzed data, conducted statistical analyses, prepared the figures, and wrote the manuscript. All authors reviewed and approved the final manuscript.

---

**Figure S1.**
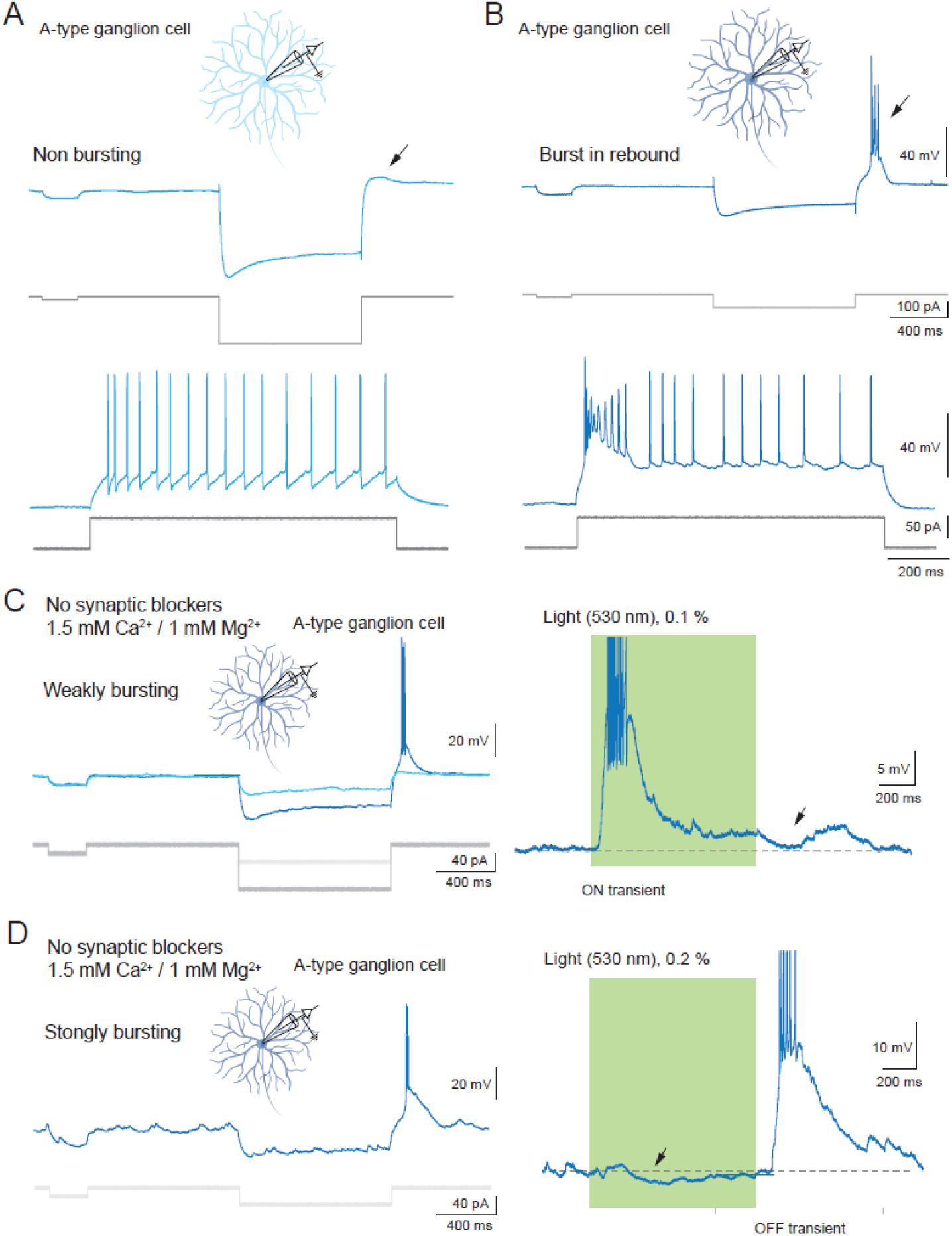
Electrophysiological and visual classification of RGCs. (A) A-type RGC without a rebound burst after strong hyperpolarization, classified as non-bursting RGC. (B) A-type RGC with a prominent rebound burst after hyperpolarization, classified as bursting RGC. (C) ON-type response in a weakly bursting RGC recorded in physiological external Ca^2+^/Mg^2+^ (1.5 / 1.0 mM). Left, electrophysiological bursting signature; right, a full-field visual stimulus evokes an excitatory ON response with spikes (truncated) and a small inhibitory OFF response (arrow). (D) OFF-type response in a strongly bursting RGC recorded in physiological external Ca^2+^/Mg^2+^ (1.5 / 1.0 mM). Left, electrophysiological bursting signature; right, a full-field visual stimulus evokes a small inhibitory ON (arrow) and a robust OFF response with spikes (truncated). RGC schematics were created with BioRender.

**Figure S2.**
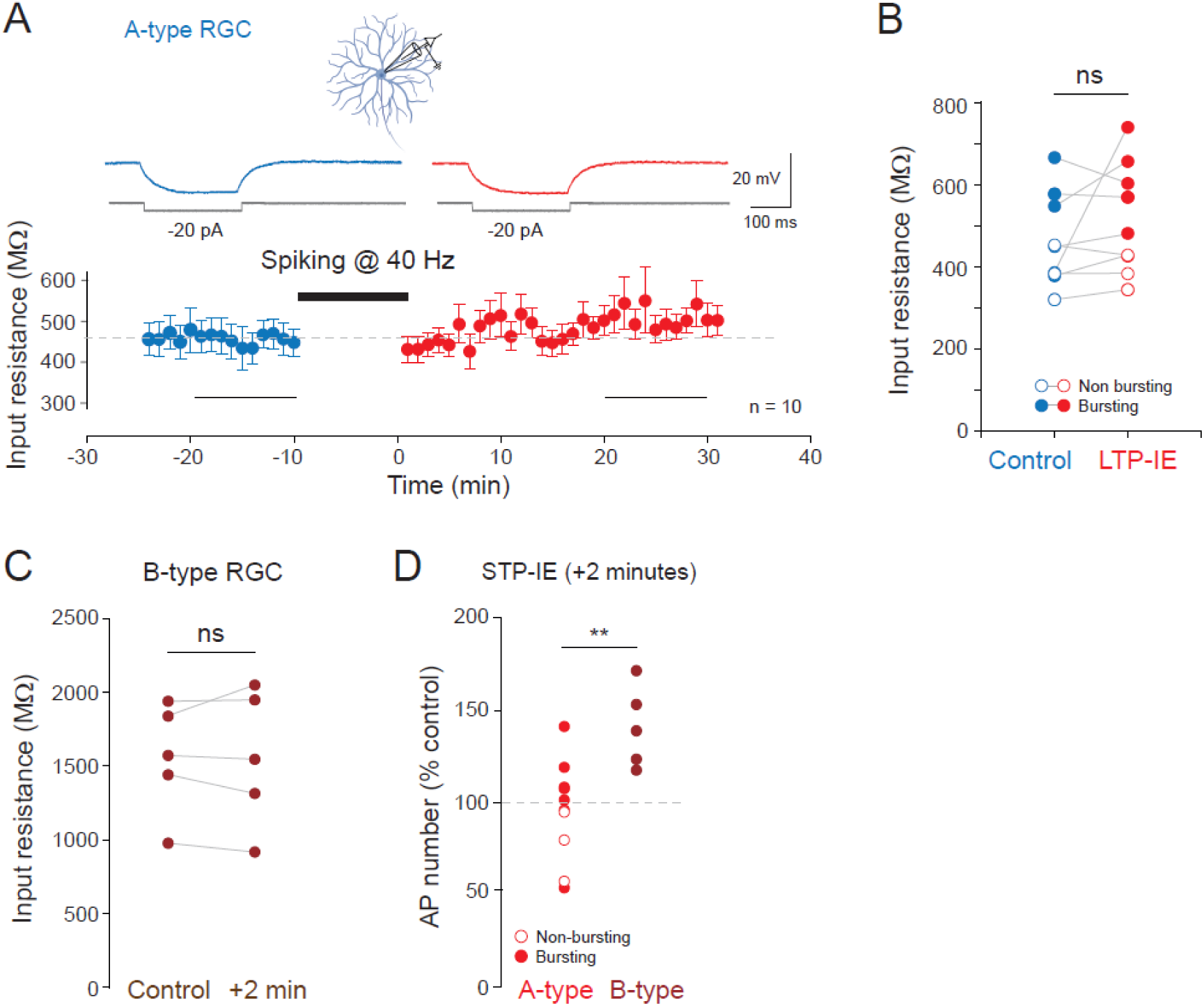
Input resistance is unchanged after intrinsic-plasticity induction in A- and B-type RGCs. (A) Time course of input resistance (Rin) in A-type RGCs before and after 40-Hz induction. (B) Paired Rin values before and after LTP-IE induction in A-type RGCs. ns, Wilcoxon signed-rank test, p > 0.1. (C) Paired Rin values in B-type RGCs before and 2 min after 40-Hz firing. ns, Wilcoxon signed-rank test, p > 0.1. (D) Normalized AP number 2 min after 40-Hz firing in A-versus B-type RGCs. **, Mann-Whitney U-test, p < 0.01. RGC schematic created with BioRender.

**Figure S3.**
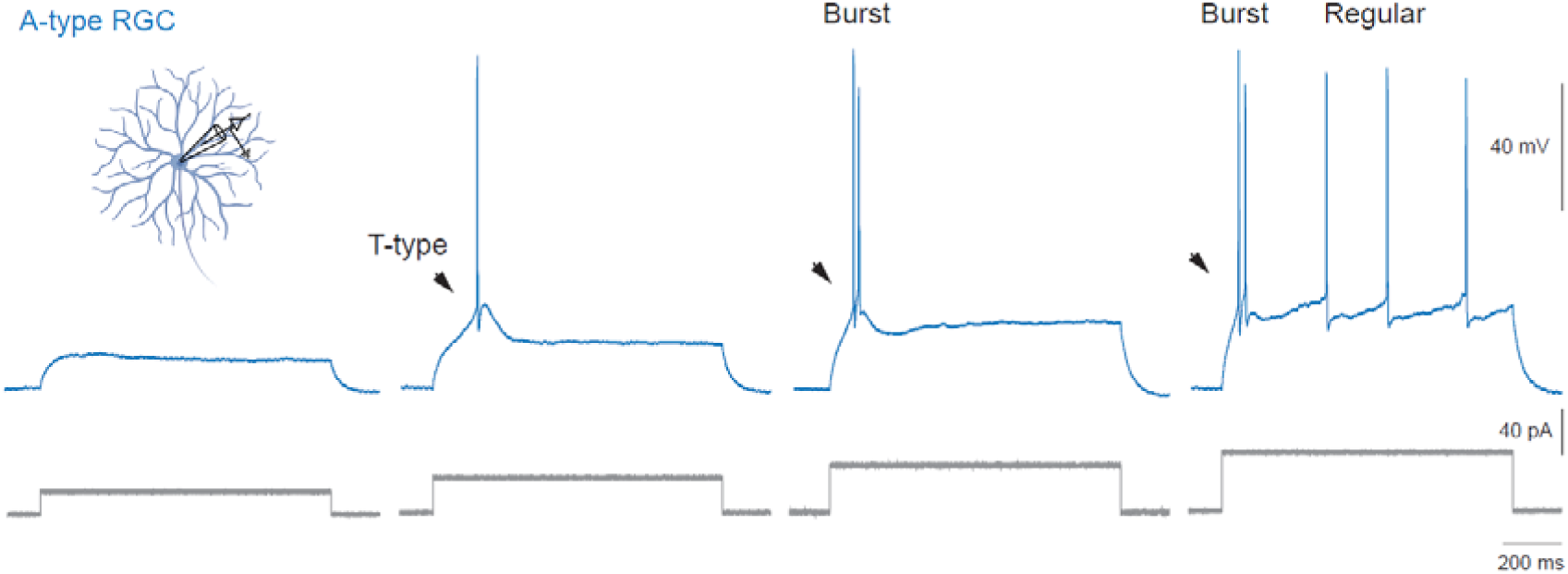
T-type calcium potential in an A-type bursting RGC. Representative voltage responses to increasing depolarizing current steps. Arrowhead, T-type calcium potential evoked by minimal depolarization from -65 mV. Larger depolarizing steps evoke a rebound burst followed by regular spiking. RGC schematic created with BioRender.

**Figure S4.**
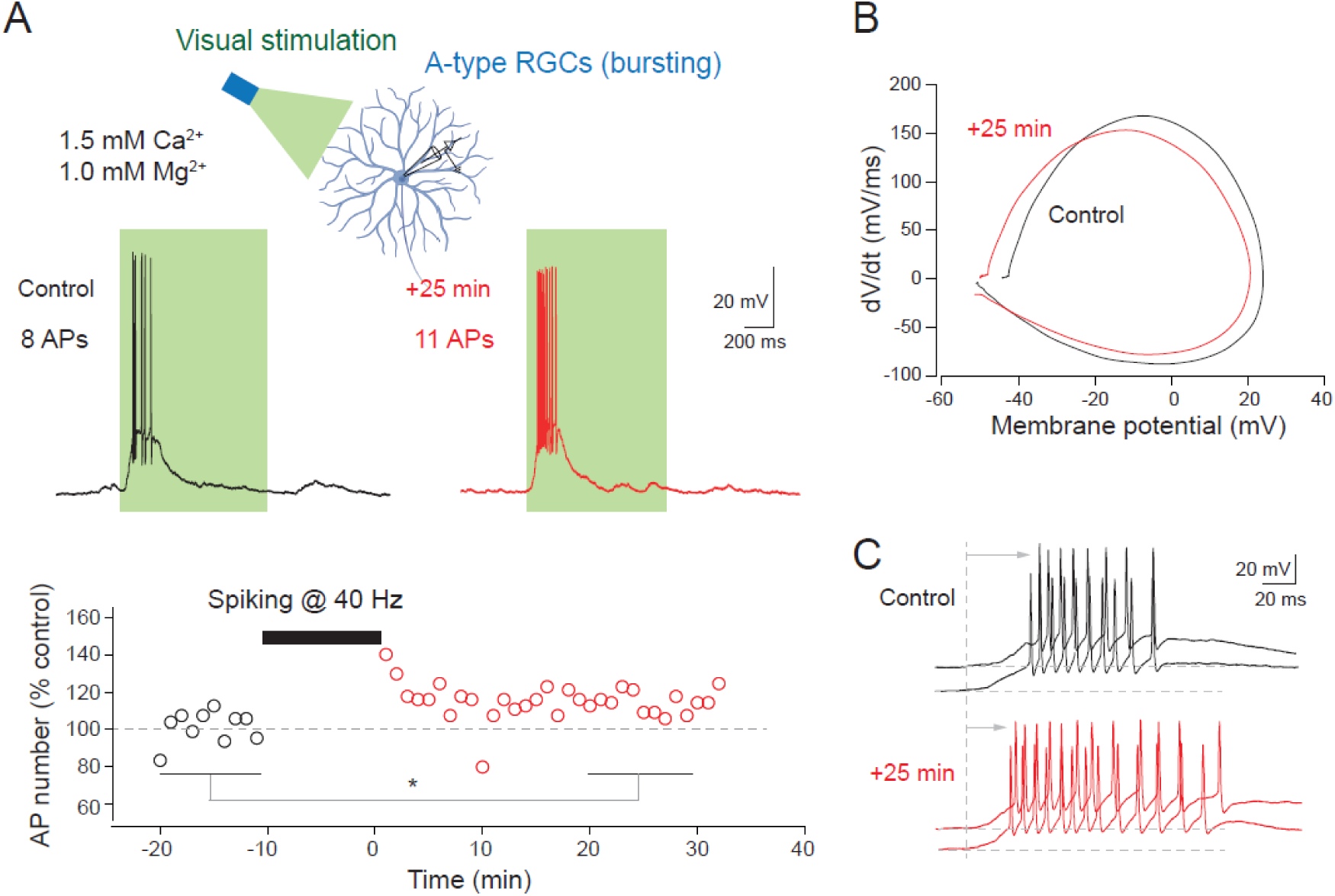
LTP-IE measured on visual responses. (A) Lasting enhancement of spiking activity evoked by green light in a A-type bursting RGC. Top, scheme of the recording configuration. Middle, representative voltage traces before (black) and after (red) spiking activity at 40 Hz showing LTP-IE of visually evoked responses. Bottom, time course of AP number before and after 40 Hz firing. *; Wilcoxon test, p<0.05. (B) Hyperpolarized AP threshold. Phase plots of the first spike before (black) and after (red) LTP-IE induction showing an ∼5 mV hyperpolarization of the AP threshold. (C) Shortening of the AP delay after LTP-IE. The vertical dashed line indicates the beginning of the visual response. RGC schematic created with BioRender.

